# Allele expression biases in mixed-ploid sugarcane accessions

**DOI:** 10.1101/2021.08.26.457296

**Authors:** Fernando Henrique Correr, Agnelo Furtado, Antonio Augusto Franco Garcia, Robert James Henry, Gabriel Rodrigues Alves Margarido

**Affiliations:** Department of Genetics, University of São Paulo, “Luiz de Queiroz” College of Agriculture, Av Pádua Dias, 11, Piracicaba 13400-970, Brazil; Queensland Alliance for Agriculture and Food Innovation, University of Queensland, Brisbane 4072, Australia

**Keywords:** Allelic imbalance, Polyploid, Allele dosage, Bayes, *Saccharum*

## Abstract

Allele-specific expression (ASE) represents differences in the magnitude of expression between alleles of the same gene. This is not straightforward for polyploids, especially autopolyploids, as knowledge about the dose of each allele is required for accurate estimation of ASE. This is the case for the genomically complex *Saccharum* species, characterized by high levels of ploidy and aneuploidy. We used a Beta-Binomial model to test for allelic imbalance in *Saccharum*, with adaptations for mixed-ploid organisms. The hierarchical Beta-Binomial model was used to test if allele expression followed the expectation based on genomic allele dosage. The highest frequencies of ASE occurred in sugarcane hybrids, suggesting a possible influence of interspecific hybridization in these genotypes. For all accessions, ASEGs were less frequent than those with balanced allelic expression. These genes were related to a broad range of processes, mostly associated with general metabolism, organelles, responses to stress and responses to stimuli. In addition, the frequency of ASEGs in high-level functional terms was similar among the genotypes, with a few genes associated with more specific biological processes. We hypothesize that ASE in *Saccharum* is largely a genotype-specific phenomenon, as a large number of ASEGs were exclusive to individual accessions.

## Introduction

Sugarcane is one of the most important polyploid crops, and is cultivated in 26.8 million hectares worldwide (FAOSTAT, 2021). Profitable and sustainable production relies on high-yielding cultivars developed by breeding programs (Scortecci *et al*., 2012). Sugarcane breeders can use molecular markers and genomic sequences to explore the variability among *Saccharum* accessions, and enhance knowledge about the molecular basis of desired traits (Mancini *et al*., 2017). However, modern cultivars are complex polyploids, which poses challenges for analyzing their genomes. Although such cultivars have a basic chromosome set of x = 10, they are highly polyploid and aneuploid interspecific hybrids, resulting in a genome of approximately 10 Gb (Diniz *et al*., 2019; Piperidis *et al*., 2010; Piperidis and D’Hont, 2020; Scortecci *et al*., 2012). These genotypes can also show a variable number of chromosomes among the homoeologous groups (Piperidis *et al*., 2010; Piperidis and D’Hont, 2020).

There are genome references recently available for hybrids and the wild species *S. spontaneum* (Garsmeur *et al*., 2018; Souza *et al*., 2019; Zhang *et al*., 2018). Although they are important genomic resources, they are still limited in the representation of multiple copies and similar alleles. In addition, multimapping is still an important problem when assigning reads to similar sequences. The hom(oe)ologous chromosomes have high levels of collinearity and gene structural conservation, as the three *Saccharum* genomes are similar (Pompidor *et al*., 2021). Association between genotypic and phenotypic data is thus not trivial in sugarcane. Instead of relying only on genomic information, approaches using transcriptomes have proven useful in investigation of likely cellular functions of putative genes, aiming to obtain molecular markers from functional genomic regions. Thus, analyses of transcriptomic data have made it possible to assess gene expression to compare different organs and developmental stages (Casu *et al*., 2007; Mattiello *et al*., 2015) and to contrast specific genotypes (Vicentini *et al*., 2015) or groups of accessions (Correr *et al*., 2020; Kasirajan *et al*., 2018).

Differential expression analysis identifies significant changes in the intensity of gene expression, revealing possible changes in metabolic pathways according to contrasting factors used in the experimental design (Soneson and Delorenzi, 2013; Wang *et al*., 2009). However, there is also variation inherent to the allelic origin of each transcript, because a heterozygous locus can have more than one haplotype being transcribed. The magnitude of variation between the expression of the haplotypes can differ, resulting in preferentially expressed alleles (Castel *et al*., 2015). Significant differences in the expression of the alleles are due to effects in proximal regulation, changes in the reading frame and epigenetic modifications (Castel *et al*., 2015). Therefore, to measure allele-specific expression (ASE), polymorphisms must be detected and the expression level of each allele be obtained via RNA sequencing (Hu *et al*., 2016; Romanel *et al*., 2015). The objective is to detect deviations from equivalent expression between the alleles (*i.e*., allelic imbalance), as well as to compare the relative allelic proportion in samples from different environments (Ereful *et al*., 2016; Tuch *et al*., 2010; Wood *et al*., 2015).

In diploids, tests for allele-specific expression often use a binomial model with an expected probability of equivalent expression between the alleles (Figure 1 - A in Additional file 1). Then, ASE stems from significant deviations from similar expression levels of both alleles. In polyploids, the allelic frequency in the homology group can influence the relative expression levels. Therefore, the doses of the alleles in each heterozygous site must be estimated for accurate assessment of ASE. The modeling procedure also follows the binomial distribution, but the expected proportion should be equivalent to the relative dose of the allele. Pham and colleagues (Pham *et al*., 2017) considered the possible dosage values in autotetraploid potato - simplex, duplex and triplex (DePristo *et al*., 2011; McKenna *et al*., 2010) - to determine the expected probability of allele counts. This is a case of studying allele-specific expression for an organism with fixed ploidy (Figure 1 - B in Additional file 1). They found from 2,180 to 5,270 genes showing preferentially expressed alleles in their experimental conditions - combination of six genotypes and two organs. Furthermore, all potato genotypes had more genes with ASE in the tuber than in the leaves, the former showing enrichment of genes coding for proteins responsible for the localization of macromolecules and transport processes. These authors emphasized that ASE reflected the breeding history of this crop, as it was more frequent in the target of selection - the tuber. On the other hand, they also reported the occurrence of ASE in genes related to traits introgressed from wild genotypes.

Polyploidy arises by whole genome duplications (WGD), originating as autopolyploids; or by hybridization between related species, resulting in allopolyploids (Lavania, 2013; Spoelhof *et al*., 2017). While the former event creates multiple sets of homologous chromosomes, the latter results in parental subgenomes that can be grouped in sets of homoeologous chromosomes (Spoelhof *et al*., 2017). The six *Saccharum* species are polyploids with a large number of chromosomes (Piperidis *et al*., 2010; Zhang *et al*., 2016). Most of the sugarcane cultivars are hybrids between *Saccharum officinarum* and *S. spontaneum*, with variable and genotype-specific numbers of chromosomes (Piperidis *et al*., 2010; Piperidis and D’Hont, 2020). Because both species are considered autopolyploids (Zhang *et al*., 2012), commercial sugarcane cultivars are interspecific hybrids that can be genomically classified as auto-allopolyploids (Pompidor *et al*., 2021; Zhang *et al*., 2016). Recently, sugarcane breeding has focused on the variability from wild accessions to explore traits for bioenergy production (da Silva, 2017). There is an interest in genes associated with important traits in sugarcane breeding - higher biomass production, resistance to diseases and tolerance to adverse environmental conditions. To that end, knowledge about gene regulation can provide useful targets for marker-association studies.

Recent research has addressed the detection of allele-specific expression in sugarcane. After determining haplotypes of particular genomic regions, a variable number of polymorphisms were found within the genes where allele expression was correlated to the dosage (De Mendonça Vilela *et al*., 2017; Sforça *et al*., 2019). Sforça and colleagues (Sforça *et al*., 2019) also reported difficulties in observing all the haplotypes of a region, inferring missing haplotypes based on expression data when possible. Another approach used the tetraploid *S. spontaneum* genome (Zhang *et al*., 2018) to investigate alleles of specific gene families (Cai *et al*., 2020). These results show that expression of alleles from genes coding for the Dof transcription family differed depending on the tissues examined, the developmental stages or hormone treatments. They also found that the *cis*-elements of the alleles of the same gene were associated with different functions. These studies pioneered research on allelic expression in sugarcane, but they focused on specific genic regions for a small group of genes. It would be informative to have a global view of the frequency of allele-specific expression in sugarcane, considering the transcriptomes of different *Saccharum* accessions.

Pham and colleagues (Pham *et al*., 2017) used a fixed ploidy of four homologs per group to detect SNPs with ASE in tetraploid potato (Figure 1 - B in Additional file 1). However, in a crop such as sugarcane, the ASE models must deal with variable ploidy levels (Garcia *et al*., 2013), respecting cytological results that demonstrate homoeology and aneuploidy (Figure 1 - C in Additional file 1) (Piperidis *et al*., 2010; Vieira *et al*., 2018). Nowadays, it is feasible to assess allele-specific expression in sugarcane by combining the expression data from RNA-Sequencing studies with the allelic dosages estimated through an appropriate pipeline for an organism with non-fixed ploidy (Pereira *et al*., 2018; Serang *et al*., 2012). Our main objective was to test for allele-specific expression using a model leveraging the doses of the alleles as prior information. Here we show the use of an adapted Beta-Binomial distribution, commonly used for diploid and polyploid organisms, to model ASE in mixed-ploid *Saccharum*. Finally, we suggest that this model can be easily applied to unravel ASE in other complex polyploid species.

## Results

### Number of polymorphisms obtained with the polyploid genotyping pipeline

A total of 63,712 polymorphic sites were identified in the *de novo* transcriptome reference, and we kept 37,902 SNPs after removing monomorphic or missing sites. By doing so, we only kept SNPs that were heterozygous in at least one of the genotypes. We also removed polymorphisms identified as indels, keeping 27,041 sites. Most of the SNPs sequenced at higher depth were dodecaploid, for all genotypes, with lower frequencies for lower ploidy levels (Figure 2 in Additional file 1). This finding is in agreement with cytological observations, as twelve is the most frequent ploidy among the homologs of *Saccharum* hybrids (Piperidis and D’Hont, 2020). Less stringent filters resulted in different distributions, with higher frequencies of hexaploid and octaploid loci. This may reflect lower accuracy for polymorphisms detected at lower depth of sequencing.

Another important observation was that the total number of SNPs was almost constant among the genotypes when no depth filter was applied (Figure 2 in Additional file 1). However, the number of heterozygous SNPs was higher in hybrids and *S. officinarum* (Table 2 in Additional file 1). When increasing the minimum depth filter, the genotypes SES205A and US85-1008 had fewer SNPs than the others. During the GBS protocol, these were the only genotypes without replication in the sequencing libraries. Furthermore, 75% of the transcripts had up to 2,665 bp, with an average of roughly four SNPs (Figure 3 in Additional file 1). It also observed that longer transcripts did not necessarily have more SNPs. This is likely explained by the inherent limitation of GBS to only detect SNPs in positions adjacent to the restriction enzyme recognition site. Overall, these figures show that the markers identified with the GBS pipeline are appropriate for genotyping and comparing different accessions.

After removing indels, missing and monomorphic sites, quantification of allele expression was performed with ASEReadCounter for 26,995 SNPs identified in 6,722 transcripts. We used the heterozygous sites in each genotype (numbers in Table 2 in Additional file 1) to test if the RNA-Seq proportion between both alleles deviated from the ratio observed in the GBS reads, indicating a likely imbalance between the alleles.

### Preferentially expressed alleles

The SNPs used to test for preferential expression were the heterozygous loci with a minimum of ten genomic reads and ten RNA-Seq reads. For all genotypes, SNPs showing ASE were the minority (Figure 1 and Table 3 in Additional file 1). Similarly, the number of genes showing allele-specific expression (ASEGs) were less frequent than non-ASEGs. No evidence of positional bias of the SNPs showing ASE was found (Figure 4 in Additional file 1) and we also found no evidence that SNPs in highly expressed genes were more likely to show ASE (Figure 5 in Additional file 1). Dissimilarity among genotypes calculated with ASE-SNPs was similar to that obtained with all loci. First, using either the relative dosage estimated with the genotypic data or the relative expression calculated from the RNA-Seq, hybrid genotypes were clustered with *S. officinarum* (Figure 6 - A and 6 - C in Additional file 1). A second cluster was formed by the *S. spontaneum* accessions. These groups were also consistent when using only SNPs classified as showing ASE (Figure 6 - B and 6 - D in Additional file 1), revealing that the occurrence of ASE may be used to estimate distances between accessions. In addition, relative expression of ASE-SNPs grouped SP80-3280 and RB72454 together, differently than what we observed when using all the evaluated SNPs.

**Figure 1:**
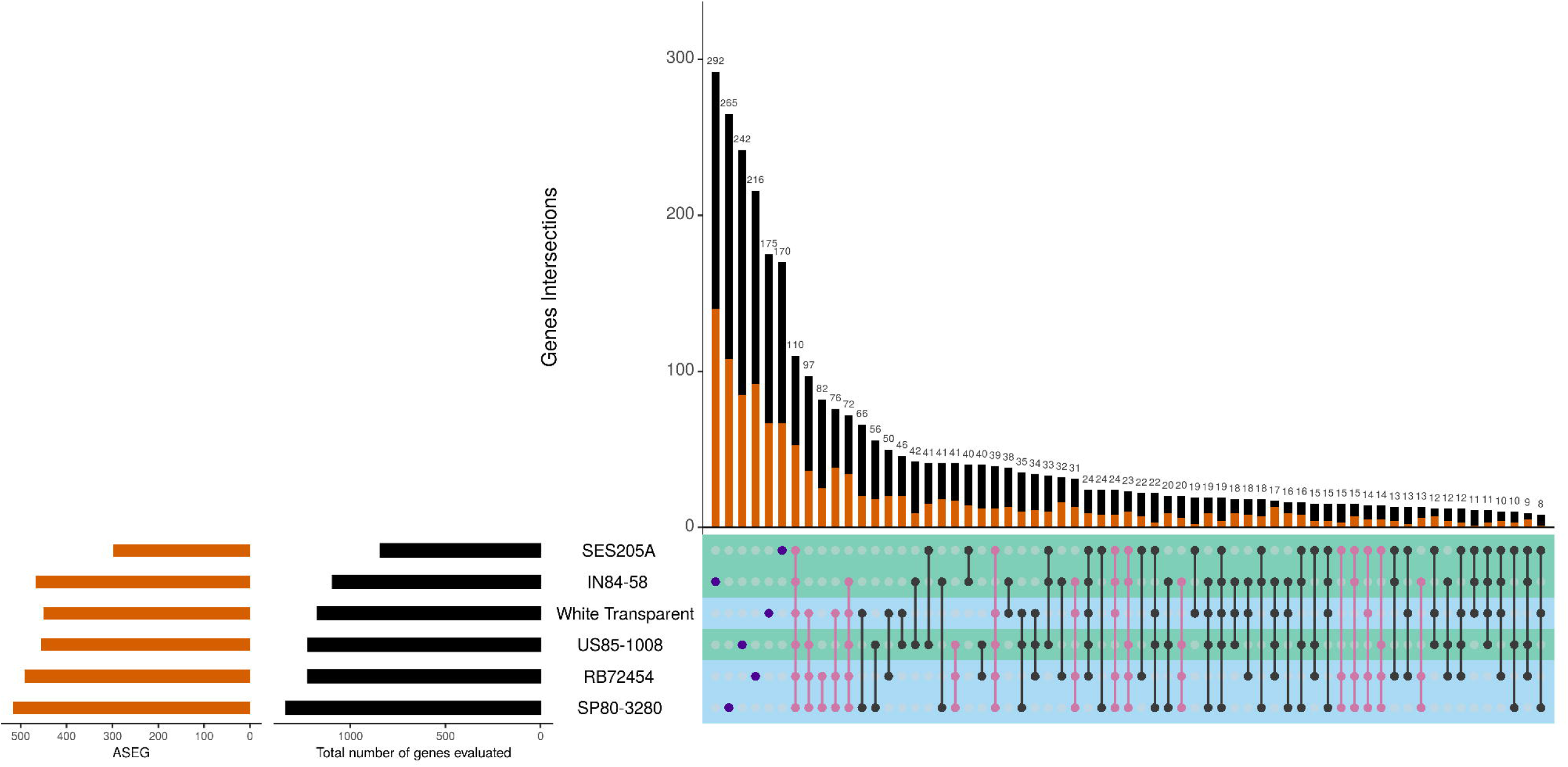
Intersections among the genes showing allele-specific expression (ASEGs) detected for each genotype. The number of ASEGs in each genotype is shown in orange and the total number of genes (ASEGs and non-ASEGs) is in black, on the left part of the plot. The right plot indicates all possible intersections among the genotypes, with ASEGs and non-ASEGs colored with the same scheme in the barplot. High fiber genotypes are shown with a green bar, and low fiber genotypes in blue. Purple dots indicate the exclusive genes of each genotype, pink dots represent the intersections where SP80-3280 and RB72454 are present.

The three hybrid genotypes - RB72454, SP80-3280 and US85-1008 - had the highest number of ASE- SNPs and the highest number of ASEGs (Figure 1). We noticed that most genes with ASE occurred exclusively in a single genotype after evaluating all possible intersections of ASEGs. However, results regarding the functional annotation were similar among genotypes (Table 4, 5, 6 and 7 in Additional file 2). We found ASEGs coding for stress-related proteins, especially disease resistance proteins. Among the disease resistance gene analogs (RGAs) and RPP genes, we found an ASEG coding for the protein *enhanced disease resistance 2*. In this gene, three SNPs revealed allele-specific expression – the two ASE-SNPs found in RB72454 were biased towards the alternative allele and one was commonly found in US85-1008 (Figure 2). The protein coded by this gene is potentially involved with hypersensitive response and in preventing senescence induced by ethylene. Curiously, a gene coding for a *probable ethylene response sensor 2* was among the ASEGs of RB72454 (Table 4 in Additional file 2).

**Figure 2:**
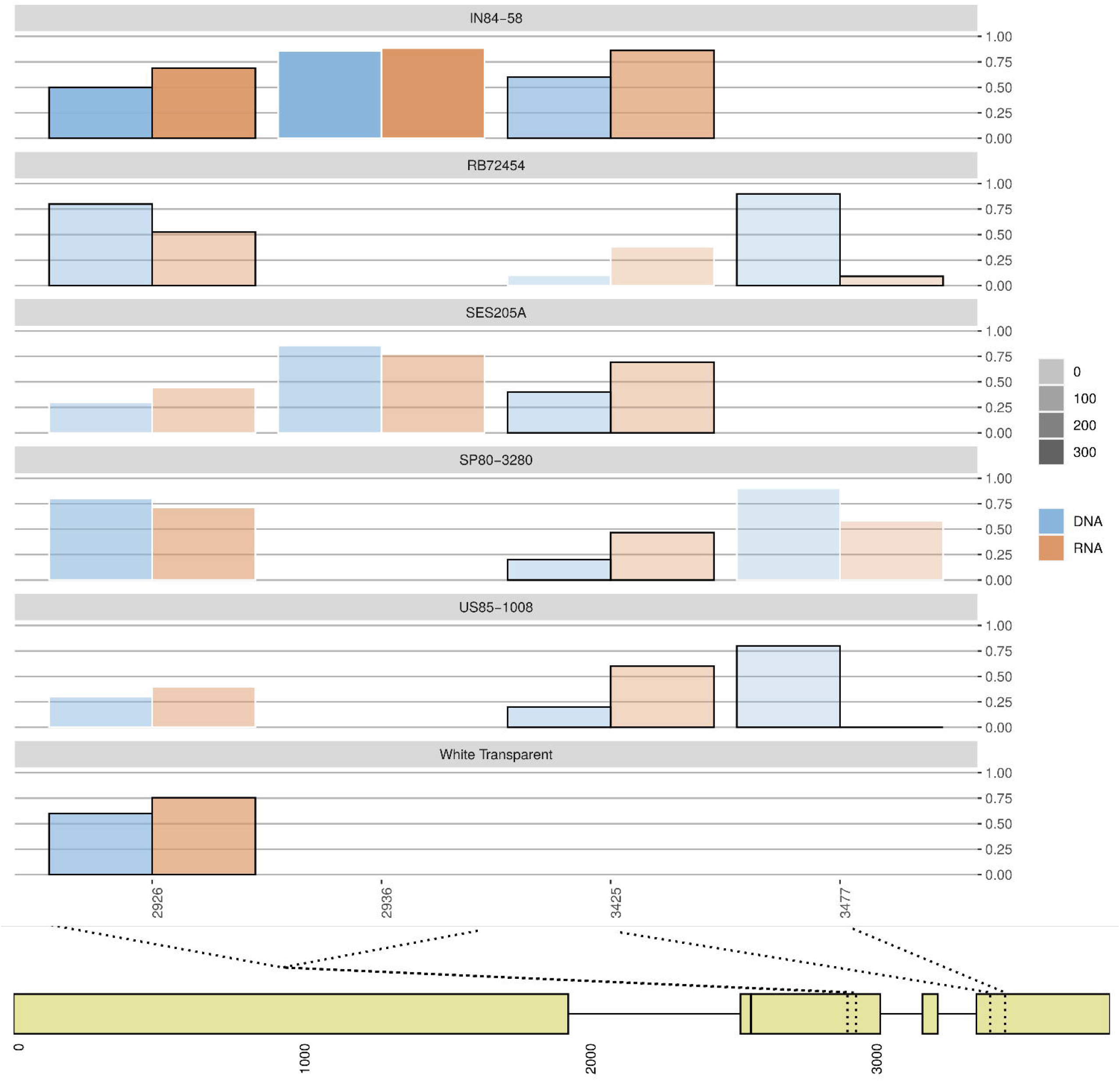
Relative genomic dose and relative expression of the reference allele from SNPs identified in the gene coding for *enhanced disease resistance 2*. Relative genomic dose of the allele is represented by a blue bar. The expressed proportion of each allele is represented by orange bars. SNPs showing significant ASE have black borders, while those not showing ASE have white borders. The color gradient represents the average expression level of the allele. The bottom part of the plot has a schematic view of the gene showing the position of the SNPs.

Sucrose content has traditionally been the focus of sugarcane breeding programs and, more recently, there has been an increasing interest in developing high fiber cultivars. Hence, ASEGs related to carbohydrate metabolism were investigated. However, a clear pattern for genes involved with carbohydrate partitioning was not identified, even in genotypes in the same phenotypic group - high or low biomass. On the other hand, genes related to this biological process were classified as ASEGs in individual genotypes. For example, a gene coding for *UTP–glucose-1-phosphate uridylyltransferase*, an enzyme involved in the synthesis of UDP-glucose, was detected. This gene had preferentially expressed alleles in all low fiber genotypes and in two high fiber accessions - IN84-58 and US85-1008 (Table 4 and 5 in Additional file 2 and Figure 3). Interestingly, the genes coding for *Sucrose-phosphate synthase* and *Sucrose transport protein SUT4*, proteins respectively involved with sucrose synthesis and transport, showed significant ASE in IN84-58. Similarly to carbohydrate metabolism, we could not find evidence of any association between photosynthesis-related ASEGs and the two phenotypic groups. Moreover, we identified genes showing genotype-specific allele-specific expression in these processes. We could identify genes for which all wild accessions had biased expression. In the case of the gene coding for *RuBisCO large subunit-binding protein*, SNPs of high-biomass genotypes and White Transparent showed preferential expression of the reference allele (Figure 7 in Additional file 1). A different pattern was observed for the ASE-SNPs found in the *phosphoenolpyruvate carboxylase 3* coding gene (Figure 8 in Additional file 1), where ASE occurred for all genotypes in at least one of the four positions.

**Figure 3:**
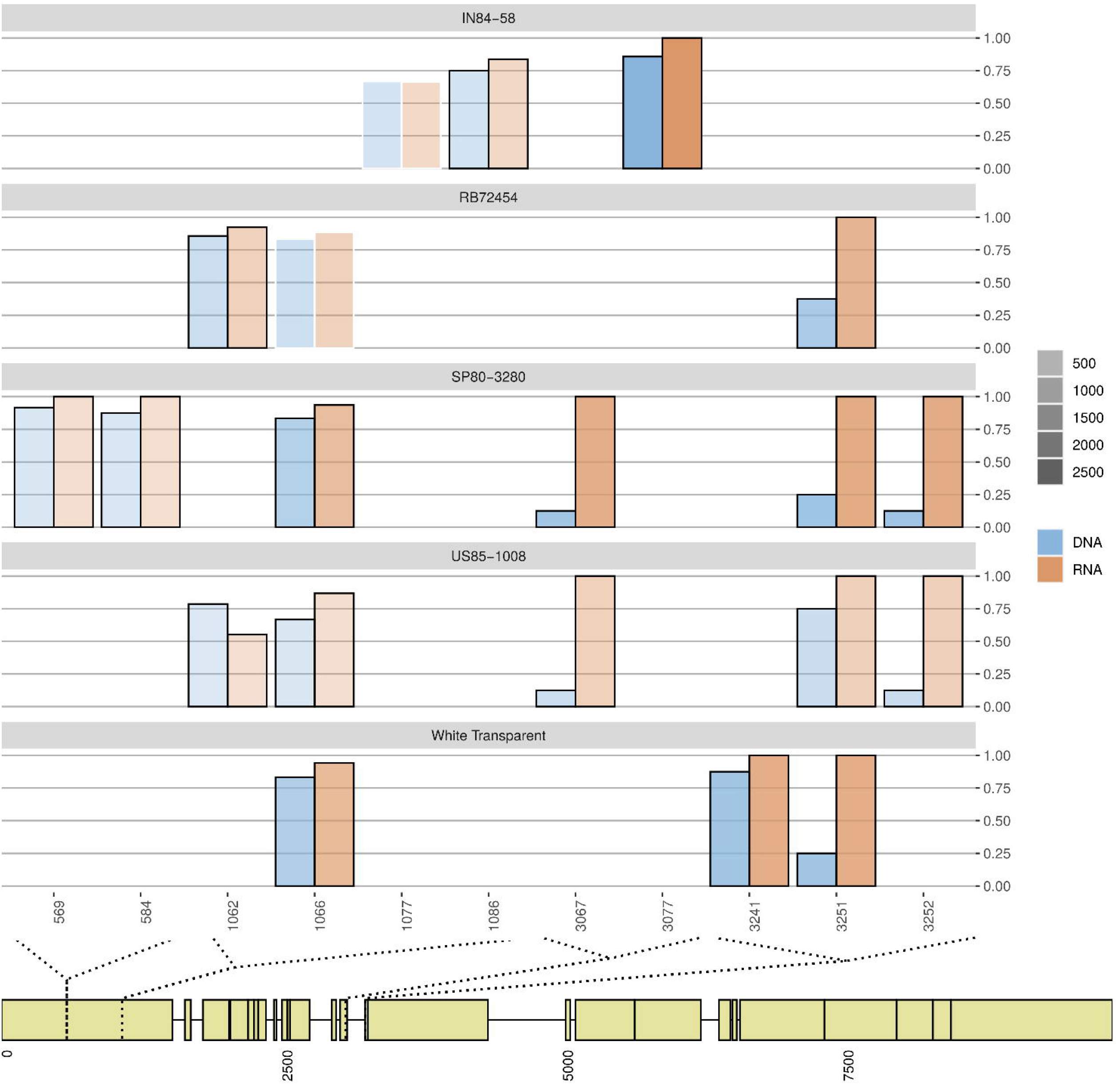
Relative genomic dosage and relative expression of the reference allele from SNPs identified in the gene coding for *UTP–glucose-1-phosphate uridylyltransferase*. Relative genomic dosage of the allele is represented by a blue bar. The expressed proportion of each allele is represented by orange bars. SNPs showing ASE have black borders, while those not showing ASE have white borders. Color gradient represents the expression level of the allele. The bottom part of the plot shows a schematic view of the transcript showing the position of the SNPs.

Functional enrichment tests were used to evaluate if ASEGs were acting on similar processes. No GO term was significantly enriched with ASEGs. This result is possibly explained by the limited number of genes with detected polymorphisms that passed the filtering steps - roughly one thousand per genotype. We then checked the frequency of ASEGs in each GO Term (Table 6 and 7 in Additional file 2) and found that GO terms with the highest frequencies of ASEGs were often found in common for all genotypes. As expected, high-level GO terms had the most ASEGs, followed by many metabolic processes and terms associated to the biosynthesis of cellular compounds. This shows that many ASEGs are possibly directly involved with maintaining the metabolism. For the GO terms, the frequency of ASEGs shared among genotypes (Table 6 in Additional file 2) is higher than the frequencies of ASEGs found in individual genotypes (Table 7 in Additional file 2). This indicates that high-level GO terms have many genotype-specific ASEGs. Hence, genes with allele-specific expression were found seemingly at random in the same pathway when considering different genotypes.

## Discussion

Da Silva (da Silva, 2017) has shown that dosage-effects and gene duplication are key factors contributing to variations in gene expression levels in sugarcane. Allele-specific expression adds another layer to the complexity of interpreting gene expression in both autopolyploids and allopolyploids. For sugarcane, this phenomenon was investigated in genes with known functions (Cai *et al*., 2020; De Mendonça Vilela *et al*., 2017). As we evaluated a larger set of expressed genes, the ASEGs found in our study were associated with a wide range of functional roles, mostly with high level metabolic processes. We found no differences among hybrids and wild genotypes regarding ASEGs related to the biosynthesis, modification, or degradation of compounds. Indeed, we observed that most ASEGs were exclusive to an individual accession (Figure 1) rather than to groups of genotypes, and that the number of ASEGs in high level GO terms was similar among the accessions. The lack of co-occurrence of ASEGs in specific pathways can be explained by two concurrent hypotheses. First, that allele-specific expression in sugarcane is genotype-specific, occurring for different genes in high level pathways. Second, there are ASEGs shared among a few genotypes that can be associated with particular functional roles. The second hypothesis could explain the few ASEGs in more specific terms (Table 7 in Additional file 2), such as the defense gene with ASE-SNPs in all genotypes (Figure 2).

Previous efforts unraveled allele-specific expression in groups of sugarcane genes. Vilela and colleagues (De Mendonça Vilela *et al*., 2017) found most of the SNPs in the TOR coding gene with the expression of different alleles matching the corresponding doses of the haplotypes. For the *Phytochrome C* coding gene, however, they identified allele-specific expression towards the main haplotype. In another endeavor, Sforça and colleagues compared the expression proportion to the genomic proportion of SNPs found in haplotypes of the genes HP600 and CENP-C (Sforça *et al*., 2019). Both genes had SNPs showing significant differences between the genomic and the transcriptomic proportions of the haplotypes. Allele expression has also been studied in combining the *S. spontaneum* genome (Zhang *et al*., 2018) and transcriptomic datasets. Recently, Cai and colleagues (Cai *et al*., 2020) used the upstream region of Dof transcription factors and found *cis*-elements associated with different functions in plants. Furthermore, these authors identified differences in upstream regions of the alleles of the same gene coding for a transcription factor. They also found alleles showing specific expression depending on tissue, developmental stage, and different hormone treatments.

These studies focused on specific haplotypes of a few genes (De Mendonça Vilela *et al*., 2017; Sforça *et al*., 2019) or evaluated specific gene families (Cai *et al*., 2020). To achieve a global view of allele-specific expression in sugarcane, we took a *de novo* transcriptome as a reference and estimated the allele dosage based on SNPs identified from GBS data. Estimating the doses is common for genotyping polyploids (de C Lara *et al*., 2019; Garcia *et al*., 2013; Gemenet *et al*., 2020; Sforça *et al*., 2019). To test for allele-specific expression in sugarcane leaves, we hypothesized that the expression of the alleles followed the allelic dosages. Our results showed that less than half of the evaluated genes had at least one SNP showing ASE (Figure 1 and Table 3 in Additional file 1). We did not verify any bias associated with ASE-SNPs (Figures 4 and 5 in Additional file 1), which also correctly clustered genotypes (Figure 6 in Additional file 1). Thus, neither a restricted coverage of polymorphisms nor differences in multiplexing apparently hampered the detection of SNPs with ASE. Moreover, our results indicate that interspecific hybridization may have caused changes in allele expression, as the highest numbers of ASE-SNPs were found in RB72454, SP80-3280 and US85-1008. On the other hand, the *S. spontaneum* accessions showed the highest proportions of ASE-SNPs. In addition, hybrids and the accession IN84-58 showed the highest number of ASEGs. Nevertheless, we note that this observation should be interpreted with caution, as the sampling of polymorphisms with GBS is limited and subject to biases (Nguyen and Lim, 2019). The small number of sampled genotypes also limits broader conclusions about the frequency of ASEGs in different genetic backgrounds.

Variation in the expression levels due to allele-dosage effects can be expected in polyploids, and this could lead to variable phenotypic effects (Osborn *et al*., 2003). Appropriate knowledge of allele dosage in polyploid organisms is required to test for allele-specific expression. ASE tests used in diploids are often based on a Binomial distribution using the null hypothesis that both alleles are expressed equally (*θ* = 0.5) (Castel *et al*., 2015; Ereful *et al*., 2016). Genotyping of organisms with a fixed ploidy is feasible (DePristo *et al*., 2011; McKenna *et al*., 2010; Pham *et al*., 2017), with markers possibly having different allelic dosages. Unfortunately, this test is not suitable for organisms with variable ploidy levels, as loci can show multiple categories of heterozytes for each ploidy level. In this scenario, cytological observations on sugarcane reveal different ploidies in the homoeologous groups (Piperidis and D’Hont, 2020), expanding the categories of allelic dosages (Garcia *et al*., 2013).

Knowledge about the complete haplotypes of the homologs/homoeologs from genomic data can improve ploidy estimation (Sforça *et al*., 2019). Using SNPs, we restricted our analysis to two alleles, although in many loci the number of alleles is probably higher. For identifying multiple alleles, we should use a haplotype-based approach, which requires a large marker density or longer sequencing reads (N’Diaye *et al*., 2017; Sehgal and Dreisigacker, 2019). However, determining the ploidy of the genomic regions for a large number of loci is still nontrivial for complex polyploids. In this scenario, the best alternative still relies on estimating the doses of alleles using molecular markers (de C Lara *et al*., 2019; Garcia *et al*., 2013; Gemenet *et al*., 2020). With this approach, knowledge of the doses has been used for constructing genetic maps and improving the performance of predictive models (de C Lara *et al*., 2019; Gemenet *et al*., 2020). In addition, this information can be used to test for allele-specific expression as done for species with fixed ploidy (Pham *et al*., 2017). For those with variable ploidy levels, models should account for the dosages of alleles in each marker. This is the scenario for our *Saccharum* dataset, in which we aimed to estimate the posterior distribution of the proportion of the reference allele. We used a Bayesian hierarchical model considering the estimated doses - obtained through genotyping - as parameters of a prior Beta distribution. Because the counts of the allele - from the expression data - follow a Binomial distribution, we modeled allele-specific expression for polyploids with a Beta-Binomial distribution (Figure 9 in Additional file 1).

Sugarcane cultivars, which are interspecific hybrids, can also show different regulation of alleles coming from different homoeologs. Unfortunately, we currently do not have enough information to identify the homoeologs but only the polymorphisms. A limitation would arise if non-identifiable duplicated genes are treated as single-copy, potentially biasing read mapping (Sforça *et al*., 2019). Lastly, as stated by Vilela and colleagues (De Mendonça Vilela *et al*., 2017), we can only speculate which mechanisms are responsible for biased expression, including the regulation of promoter regions or epigenetic changes (Castel *et al*., 2015). For a deeper investigation of the causes of ASE, multiple omics approaches should be integrated. Through genomics, assessment of the upstream and downstream regions can reveal polymorphisms in *cis*-elements. These regions can be also investigated for epigenetic modifications affecting gene regulation. In any case, by combining genomic and transcriptomic data we can identify ASEGs independently from the underlying causes.

Testing for allele-specific expression is relevant to understand differences in tissues, conditions or genotypes. Previous studies in plants emphasize how this phenomenon is common among the expressed genes. Allelic-specific expression was found in more than 50% of the genes in the maize ear of a hybrid cultivar, with a similar number of ASEGs found independently of the developmental stage (Hu *et al*., 2016). They also found a higher contribution of the alleles from one parent, but this was less pronounced during floret differentiation. Ereful and colleagues (Ereful *et al*., 2016) studied allelic imbalance combining rice genotypes (parents and F1 hybrids) and drought conditions (plants under normal water regime or following a dry-down protocol). They suggested that the occurrence of ASE was more associated to the genotype than due to water stress. However, depending on the crop, more ASEGs can be found in specific tissues. Pham and colleagues (Pham *et al*., 2017) found evidence that allele-specific expression is more frequent in potato tubers than in leaves, probably due to the selection for carbohydrate accumulation in the tubers.

Allopolyploids - or even a diploid interspecific hybrid - can be compared to their diploid parents to verify the occurrence of expression level dominance and also homoeolog-specific expression (Yoo *et al*., 2013). The study of alleles in allopolyploids relies on assessing the expression of the homoeologs to test for both expression level dominance and homoeolog-specific expression. This depends on previous knowledge of the allopolyploid parents, as the aim is to verify possible biases in gene expression towards a subgenome (Grover *et al*., 2012). The analysis is similar to that performed in the maize hybrid by Hu and colleagues (Hu *et al*., 2016) to determine the parental alleles with biased expression. In cotton, transgressive expression and expression level dominance of the A or D genome, together, were more frequent than additive expression (Yoo *et al*., 2013). However, homoeolog-specific expression was balanced between the subgenomes, which was partially explained by differential regulation of one parental homoeolog despite the expression level dominance of the other parental genome. In addition, allele-specific expression of polyploids can be more associated with the genotype than to other factors. Similarly, Powell and colleagues (Powell *et al*., 2017) stated that homoeolog expression bias was inherent to the wheat genotype, while the infection by necrotrophic *Fusarium pseudograminearum* mostly altered the magnitude of expression of the subgenomes. Knowledge of gene expression in the parental subgenomes is still lacking in the literature for studying expression level dominance and homoeolog-specific expression in sugarcane.

Genes showing preferential allele expression can be used for targeted genotyping to discover QTL regions associated with a trait. ASEGs found in rice subjected to different drought treatments were closely located to eight markers surrounding QTLs with effects on grain yield under drought (Ereful *et al*., 2016). When implemented in breeding of polyploid crops, estimation of doses can improve phenotypic predictions compared to the diploid approximation for heterozygous loci. According to De Lara and colleagues (de C Lara *et al*., 2019), the predictive ability of genomic selection models was higher when considering allele dosages in the autotetraploid *Panicum maximum*. In addition to using genomic doses, knowledge of expression biases could improve the accuracy of predictive models for plant breeding, especially in the genomic selection context. Depending on the trait evaluated, ASEGs are potential targets for associating genomic regions and phenotypes. Nowadays, sugarcane breeding focuses on bioenergy-associated traits (da Silva, 2017). Assessing the regulation of allele expression in *Saccharum* can provide targets to help in this process.

Allele-specific expression of a large set of expressed genes has been evaluated in plants (Ereful *et al*., 2016; Hu *et al*., 2016), but these studies are still limited in polyploids (Pham *et al*., 2017; Powell *et al*., 2017; Yoo *et al*., 2013). Polyploidy has a significant role in the evolution of plants and many crops are recognizable polyploids, while others experienced ancient polyploidization (Osborn *et al*., 2003). Among the most important polyploid crops, sugarcane presents a complex genome, with a large set of chromosomes. Indeed, the wild species used to develop modern cultivars, *S. officinarum* and *S. spontaneum*, are polyploids showing up to ten groups of homologs, at least six chromosomes per group and aneuploidy is frequent in some accessions (Piperidis and D’Hont, 2020). For this reason, we aimed to shed light on the occurrence of allele-specific expression in the genus by assessing a set of wild and hybrid genotypes. In this report, we aimed to assess allelespecific expression in sugarcane using a large set of genes and multiple genotypes. To achieve this objective, we modified the commonly used Beta-Binomial model to appropriately assess allele-specific expression in mixed-ploid organisms. This model can be easily applied to other polyploids, both with fixed and variable ploidy levels.

## Material and Methods

### Biological material, SNP calling pipeline and quantification of allele expression

Genotypic and transcriptomic information of *Saccharum* genotypes was used to investigate the expression of different alleles. To characterize expression profiles, we used a set of six genotypes with three replicas each from a previous gene expression study (Correr *et al*., 2020) - IN84-58, RB72454, SES205A, SP80-3280, US85-1008 and White Transparent. These genotypes represent two groups of accessions contrasting in key biomass traits - fiber content and tillering capacity. Genotypes of the high biomass group include the hybrid US85-1008 and the *S. spontaneum* genotypes IN84-58 and SES205. The low biomass group included the *S. officinarum* White Transparent and the hybrids RB72454 and SP80-3280. Briefly, we collected portions of the first visible dewlap leaves (+1) from six-month-old sugarcane plants and extracted the total RNA from the middle section of each leaf. Pooled libraries were sequenced in two lanes of an Illumina HiSeq 2500 platform, in paired-end mode (2 × 100 bp). Information regarding those genotypes can be found in the supplementary material (Table 1 in Additional File 1). Herein we used as a reference the longest isoforms of a transcriptome assembled *de novo* using the RNA-Seq reads of the full set of genotypes (Correr *et al*., 2020).

A panel of *Saccharum* accessions forms the Brazilian Panel of Sugarcane Genotypes (BPSG), which is composed of wild accessions and hybrids from Brazilian and foreign breeding programs (Medeiros *et al*., 2020). These accessions were genotyped using the genotype-by-sequencing (GBS) protocol (Elshire *et al*., 2011) with the PstI restriction enzyme. Library preparation was planned to provide a higher sequencing depth for some genotypes, by including duplicate samples in multiple library plates, including White Transparent, IN8458, RB72454 and, in particular, SP80-3280. A pipeline for SNP discovery in polyploids was performed with Tassel4-Poly (v.4.3.7 - modified) (Pereira *et al*., 2018), using Bowtie2 (v.2.3.3) (Langmead and Salzberg, 2012) to align the GBS reads. First, for SNP discovery we used the standard Tassel4-Poly pipeline with the following main modifications: a minimum minor allele frequency of 0.01 (*mnMAF*) and a minimum minor allele count (*mnMAC*) of 40. Next, the ploidy and allelic dosages for each site were estimated with SuperMASSA (Serang *et al*., 2012) and VCF2SM (Pereira *et al*., 2018). We used the Hardy-Weinberg inference model with a minimum call rate of 50%, a naïve posterior threshold of 0.5 and a minimum posterior probability to keep a variant of 0.5. Ploidy levels ranging from four to 16 were tested, then filtered for polymorphic sites with the most likely ploidy being between six and 14. The SNP calling process took into account all genotypes from the BPSG, but only those present in our RNA-Seq data were kept for downstream analysis. The VCF file was filtered to remove sites where the genotypes were homozygous or had missing calls, as well as those identified as insertions or deletions.

HISAT2 (v.2.1.0) (Kim *et al*., 2015) was used to align the RNA-Seq reads to the *de novo* transcriptome. Quantification of read counts of each allele was performed with the GATK ASEReadCounter tool (v.4.1.4.1) (Castel *et al*., 2015; DePristo *et al*., 2011; McKenna *et al*., 2010) for each aligned library. Counts of the reference allele and the total counts for each SNP for each genotype were scored. Reads from both lanes of the same sample were grouped, as no batch effect was identified. Sites with at least ten genomic reads were retained and positions showing low expression – less than ten RNA-Seq reads – were removed.

### Model to test for allele-specific expression in *Saccharum*

To assess the occurrence of allelic imbalance in a given SNP, we tested if the expression of the reference allele was equal to its relative dosage in the genome, given the estimated ploidy. For the *i*-th SNP of genotype *k*, *α_ik_* and *β_ik_* were the dosage of the reference and the alternative alleles, respectively (Figure 9 - Genotyping Additional file 1). First, the genomic ratio was calculated as the dosage of the reference allele divided by the corresponding ploidy level 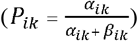. Next, the proportion of the reference allele was estimated from the RNA-Seq count, denoted by *θ_ik_*. Then, we tested the null hypothesis of no significant difference between these two ratios:

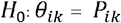

A model following the Beta-Binomial distribution was used to test this hypothesis (Figure 9 - I, II and III in Additional file 1). First, we modeled the number of RNA-Seq reads of the reference allele of the *i*-th SNP, on the *r*-th replicate of the *k*-th genotype, denoted by *y_irk_*, following a Binomial distribution (Figure 9 - II in Additional file 1):

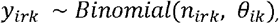

where *n_irk_* represents the total number of reads of the SNP for the corresponding sample. The prior distribution of the parameter *θ_ik_* was modeled by a Beta distribution (Figure 2 - I in Additional file 1), using as parameters the dosages of the alleles:

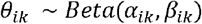

We obtained the highest density interval (HDI) of the posterior distribution of *θ_ik_* via an optimization algorithm of the Beta-Binomial distribution. We deemed a SNP as showing a preferentially expressed allele if the ratio *P_ik_* was outside of the highest HDI of *θ_ik_* (Figure 2 - III in Additional file 1). We applied a Bonferroni-like correction, by making the credibility mass for obtaining the HDI corresponding to 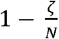, where *ζ* is the desired significance level (0.05) and *N* is the number of genes with at least one tested SNP. Although we found thousands of SNPs in each genotype (Table 2 in Additional file 1), GBS relies on the presence of enzyme restriction sites and the genotyping resulted in many adjacent and redundant SNPs. Then, because we assessed roughly a thousand independent heterozygous SNPs in each genotype – a number similar to the number of genes assessed in each genotype -, we used *N* = 1000. A gene with at least one SNP with allelic-specific expression was called as having ASE (ASEG).

### Enrichment analysis

Gene Ontology (GO) terms was evaluated for enrichment with ASEGs. To that end we used the ASEGs as the set of selected genes, compared against the background of all the genes with at least one heterozygous SNP. Tests were performed with the GOSEQ R package (v.1.38.0 in R 3.6.0) (Young *et al*., 2010). Terms with an FDR-adjusted p-value less than 5% (Benjamini and Hochberg, 1995) were considered overrepresented.

## Acknowledgements

All computational infrastructure was supported by grant #2015/22993–7, São Paulo Research Foundation (FAPESP), awarded to GRAM, and by The University of Queensland’s Research Computing Centre (RCC). This study was financed in part by the Coordenação de Aperfeiçoamento de Pessoal de Nível Superior - Brasil (CAPES) - Finance Code 001. FHC and GRAM received fellowship grants from the Institutional Program for Internationalization financed by CAPES – processes 88887.367965/2019-00 and 88887.466432/2019-00, respectively. FHC received a fellowship grant from the Brazilian National Council for Scientific and Technological Development (CNPq).

## Data Statement

The raw Illumina RNA-sequencing data used in this article are available at DDBJ/EMBL/GenBank under the BioProject ID PRJEB38368. The genotyping file of the accessions is available at Dataverse (https://dataverse.harvard.edu/privateurl.xhtml?token=b7586338-eeb7-4039-a083-0657da7114fb).

## Supporting Information

Additional File 1 – Supplementary figures 1-9 and Supplementary tables 1-3.

Additional File 2 – Supplementary tables 4-7.

