## Additional file 1 for "Allele expression biases in mixed-ploid sugarcane accessions"

##### Supplementary figure

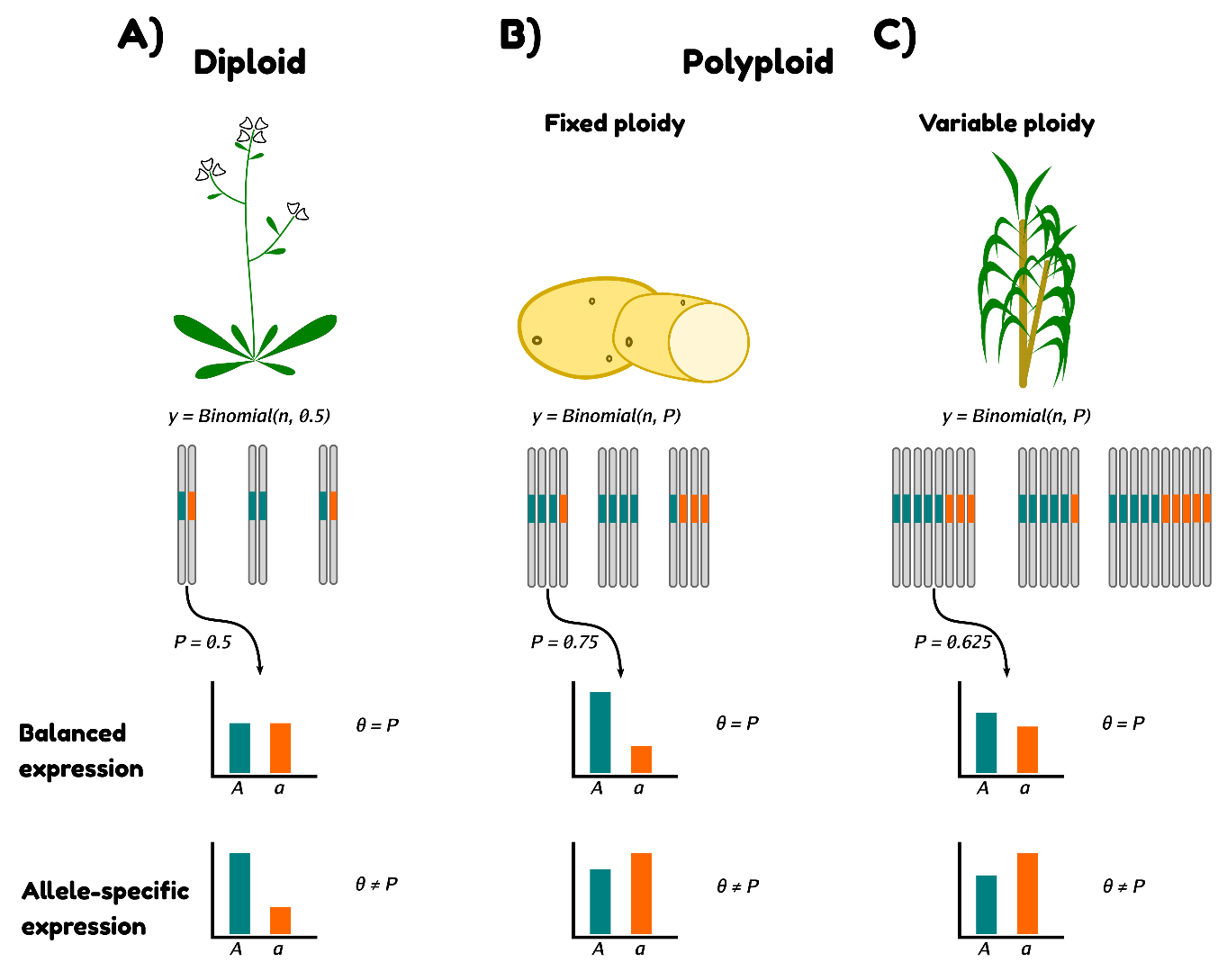

**Figure 1: Allele-specific expression studies in different ploidy scenarios.** Plants must be properly genotyped to identify homozygous and heterozygous loci. In diploid plant species, tests for allele-specific expression use a known probability of 0.5 of sampling reads with the reference allele (A). For polyploids, we rely on knowledge of the doses of each allele to calculate the proportion *P*. If the ploidy is fixed - the same in all homology groups -, *P* changes according to the doses (B). In polyploids with a variable number of homologous chromosomes per group, we need to properly estimate the ploidy of each group and use the allele doses to calculate *P* (C). If the proportion of the reference allele from the RNA-Sequencing (*θ*) is significantly different from *P*, the gene is said to show allele-specific expression.

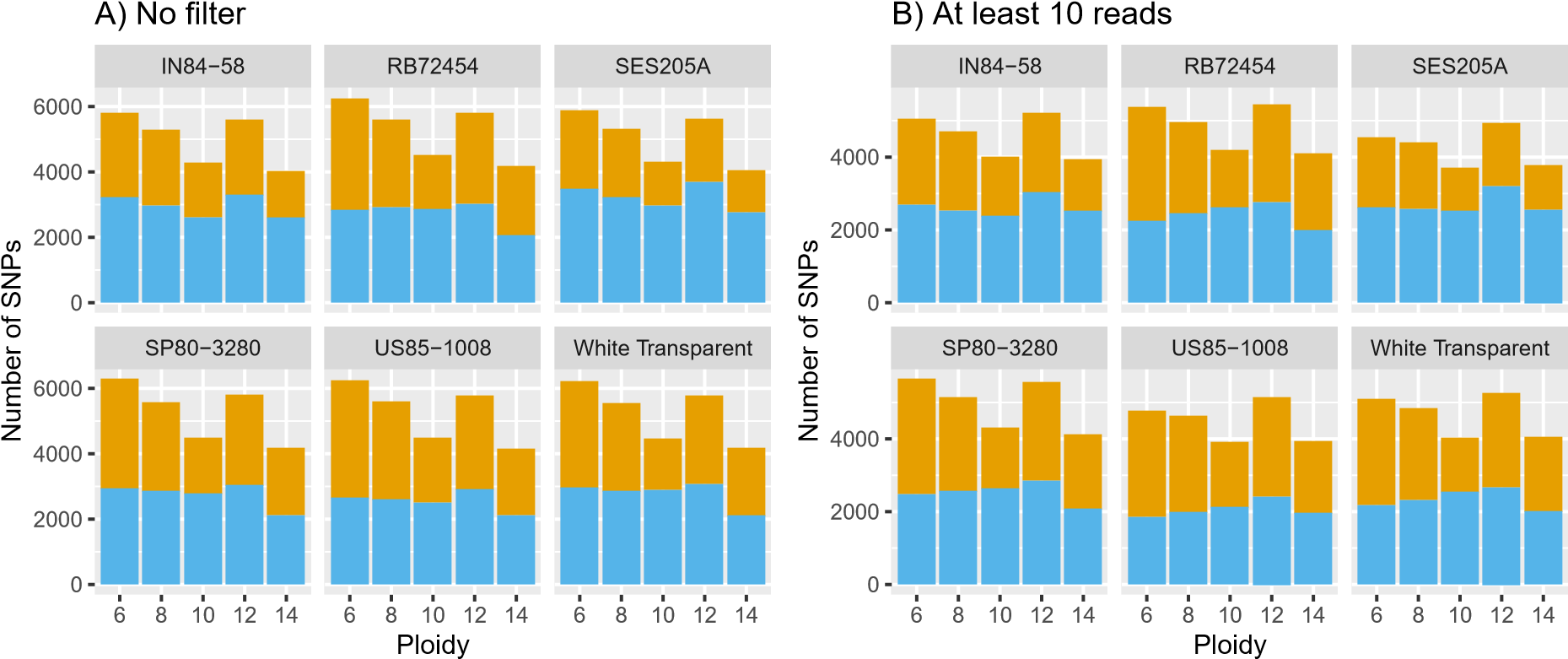

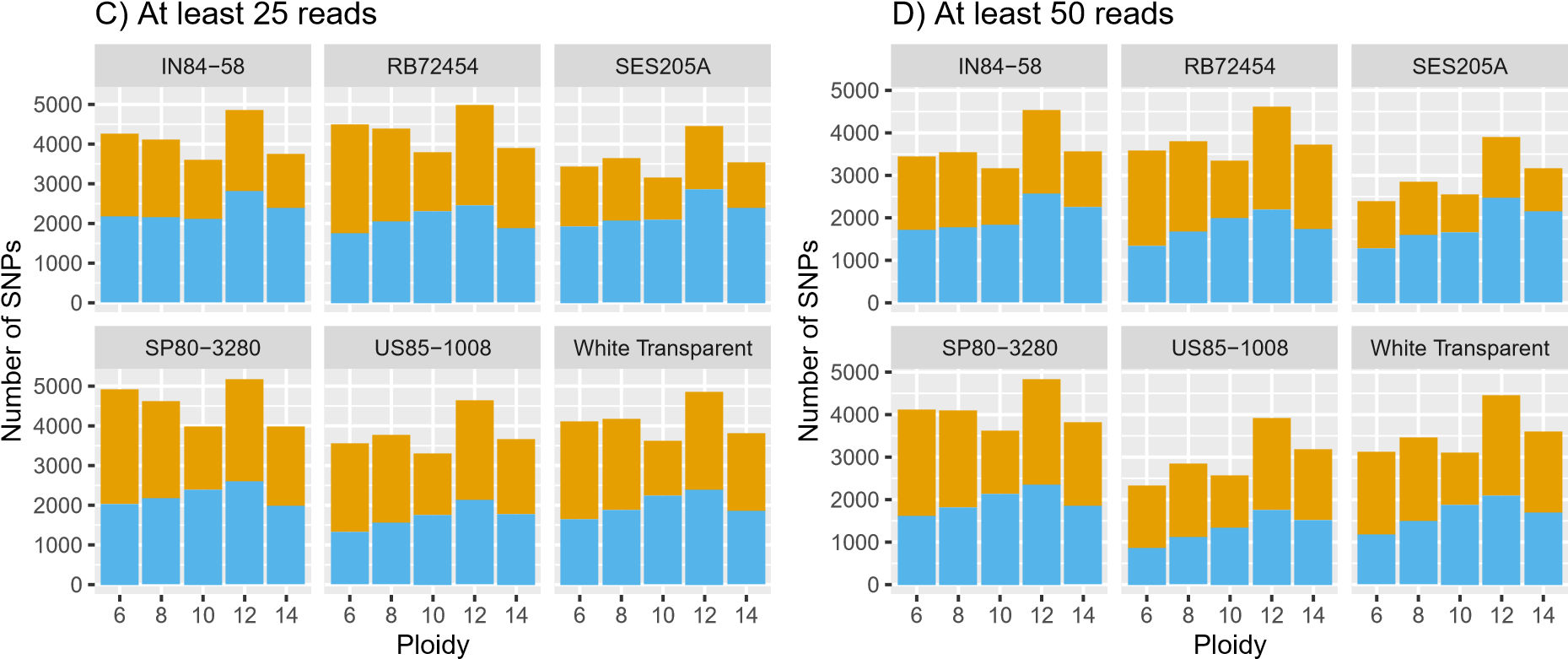

E) At least 100 reads

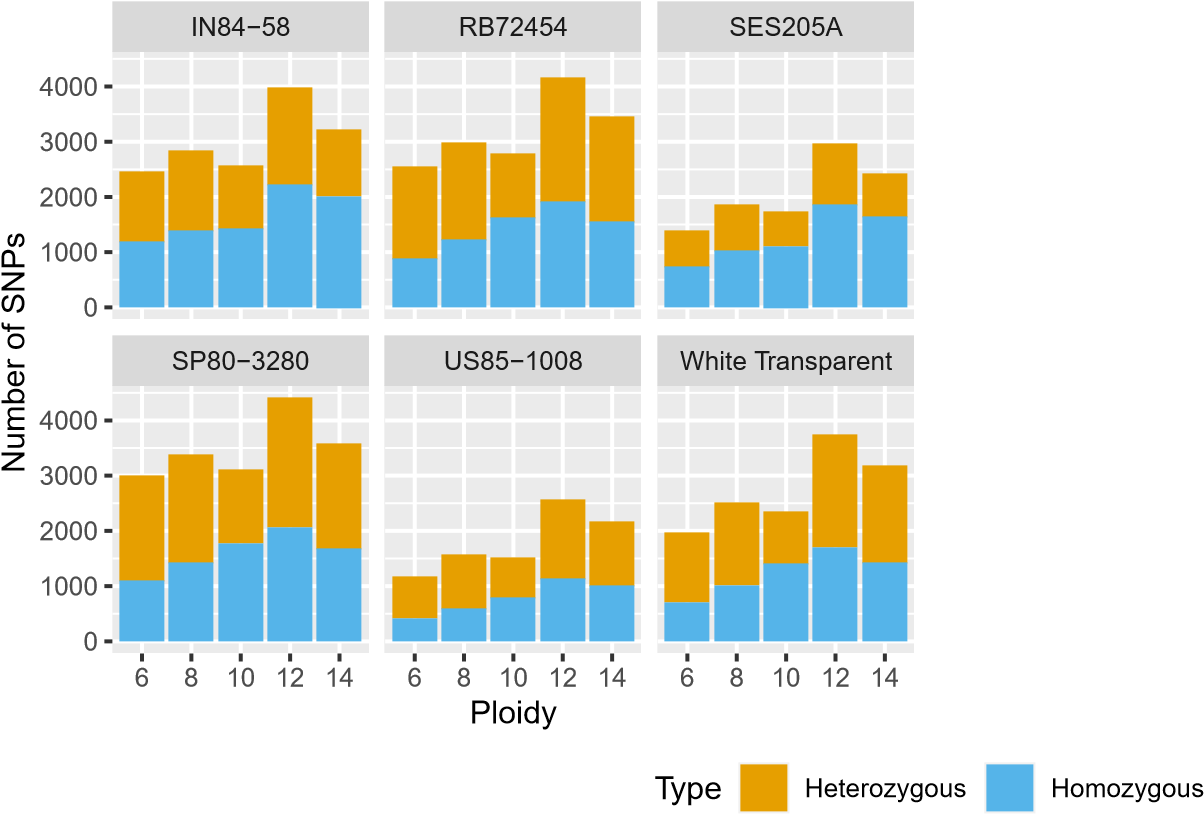

**Figure 2: Number of SNPs according to the ploidy levels for each genotype**. The number of homozygous and heterozygous SNPs for each ploidy level are presented in different scenarios: without any filter (A); with a minimum count of 10 (B), 25 (C), 50 (D) and 100 (E) GBS reads. Heterozygous SNPs are shown in orange, while the homozygous ones are in blue. Each subplot identifies a different genotype.

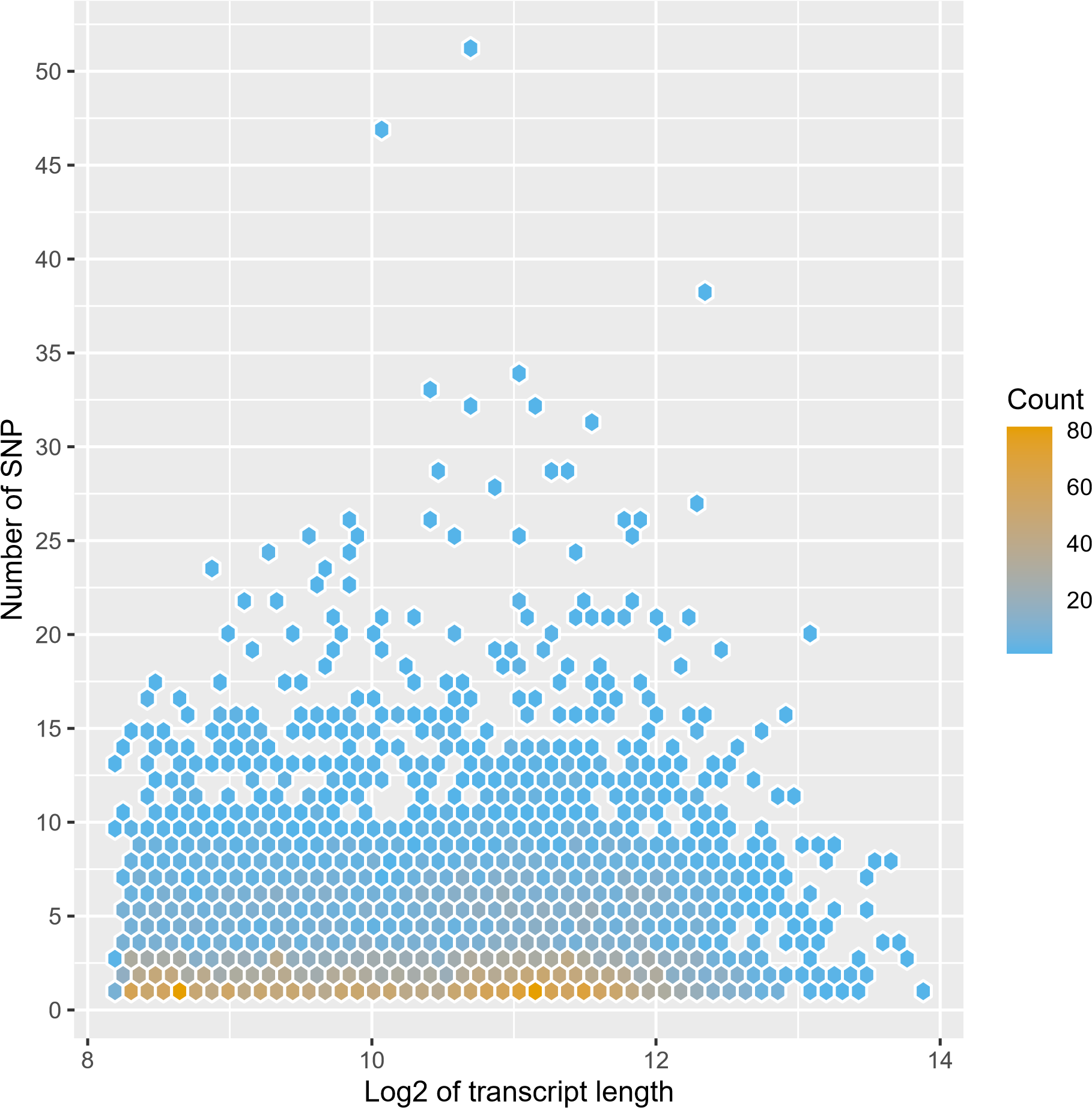

**Figure 3: Number of SNPs as a function of transcript length**. Counts are represented by a gradient from low (blue) to high numbers (orange).

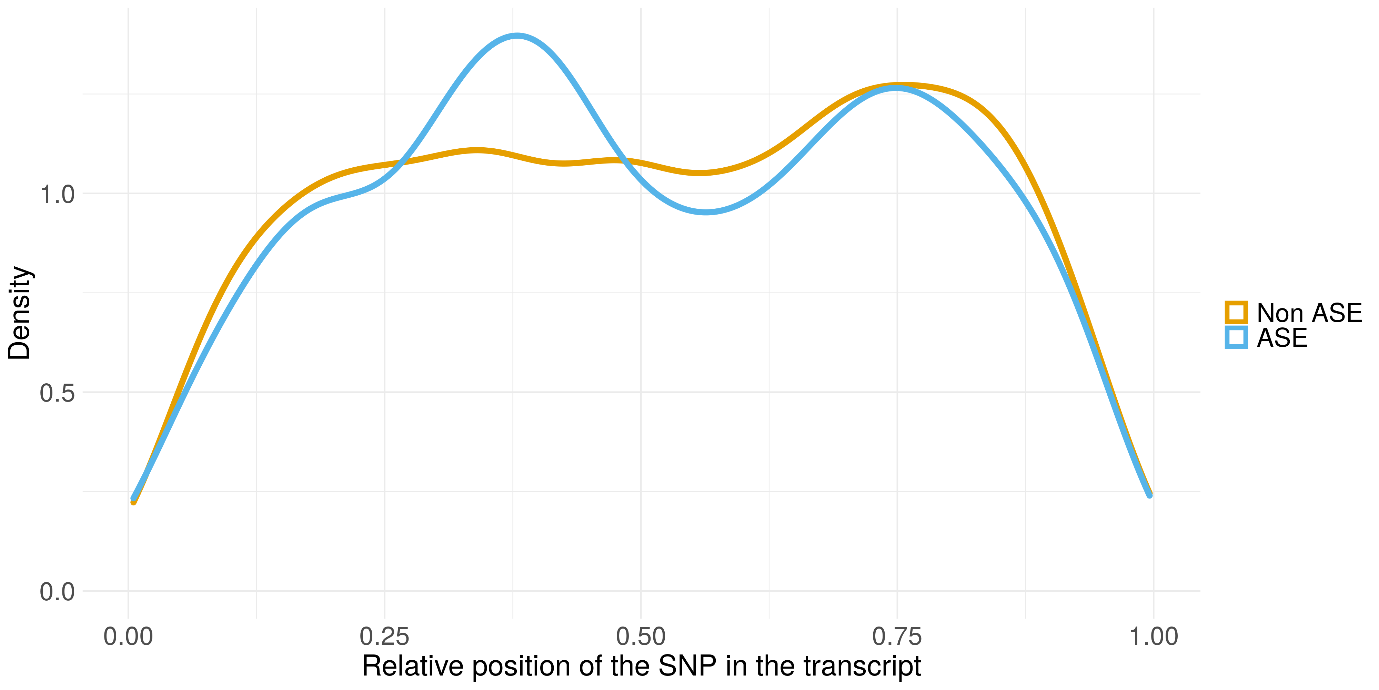

**Figure 4: Distribution of SNPs along the normalized length of transcripts**. SNPs with no allele-specific expression (ASE) are shown in orange and those with significant ASE are in blue.

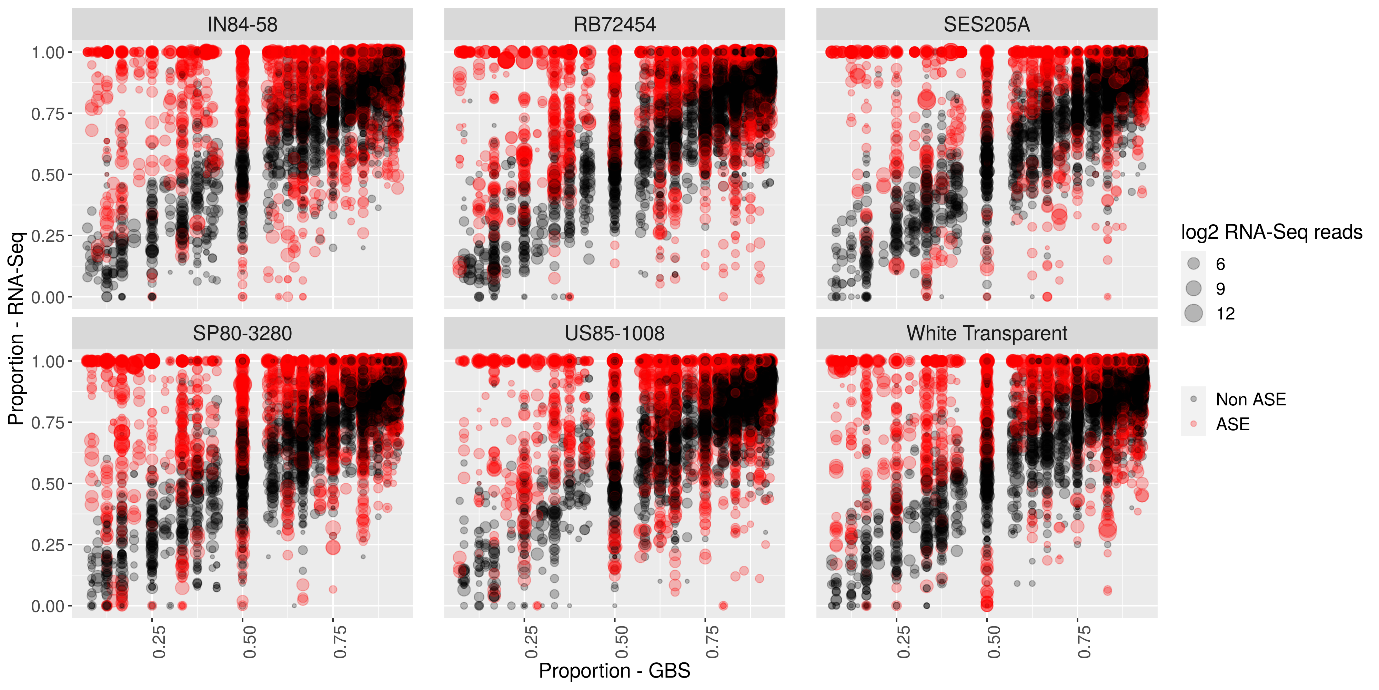

**Figure 5: Relationship between the proportion of reads with the reference allele in genomic and expression datasets**. SNPs with significant ASE are colored in red and non-ASE SNPs are colored in black. The size of each point is proportional to the overall expression level of both alleles of each SNP. Data for each genotype is shown in a different subplot.

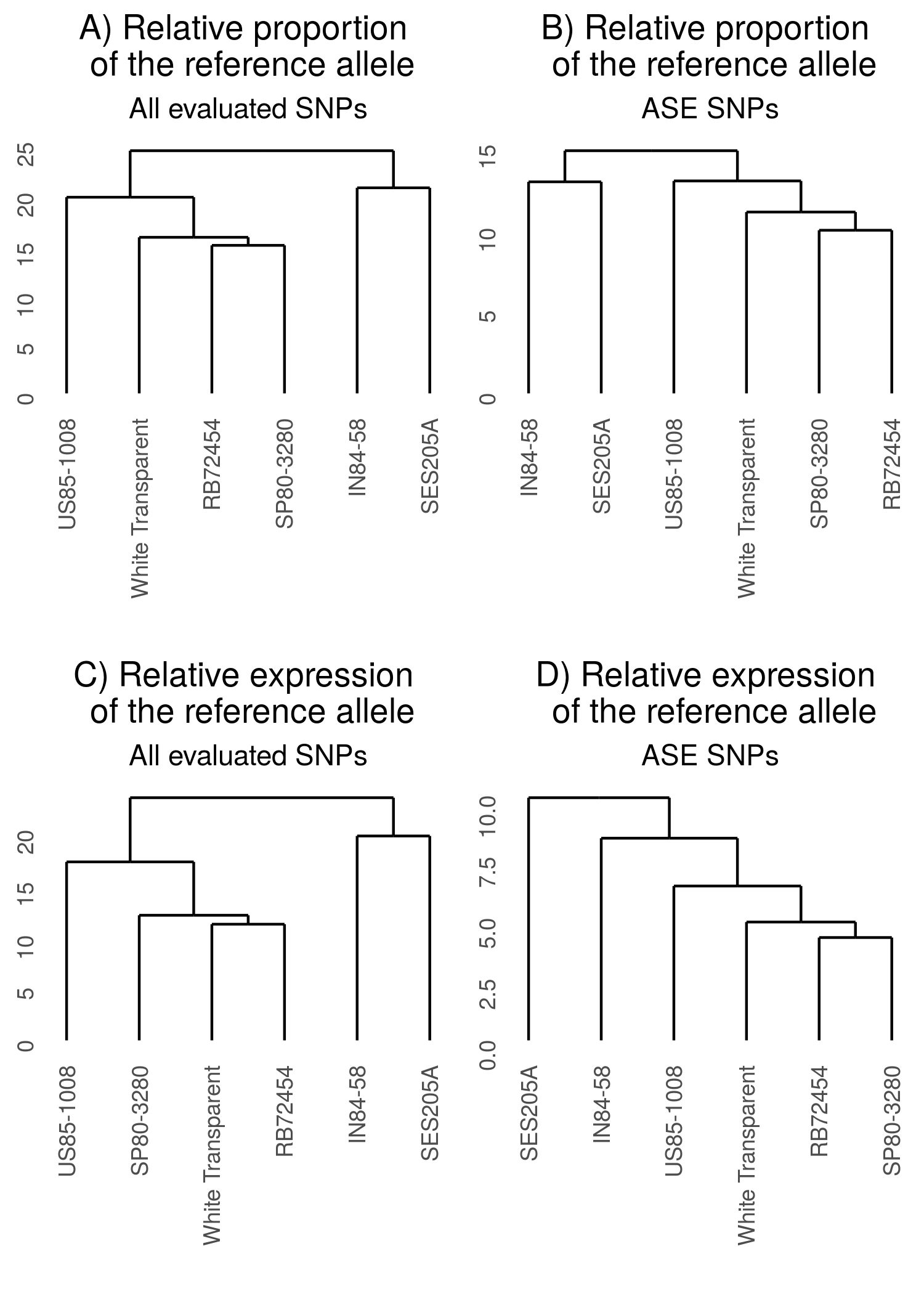

**Figure 6: Dendrograms of the hierarchical clustering of genotypes based on heterozygous SNPs**. Genotypes were clustered based on the genomic proportion of the reference allele using all heterozygous SNPs (A) and SNPs with allele-specific expression (ASE) only (B). The relative expression of the reference allele from all heterozygous SNPs (C) and ASE SNPs (D) was also used to cluster the genotypes.

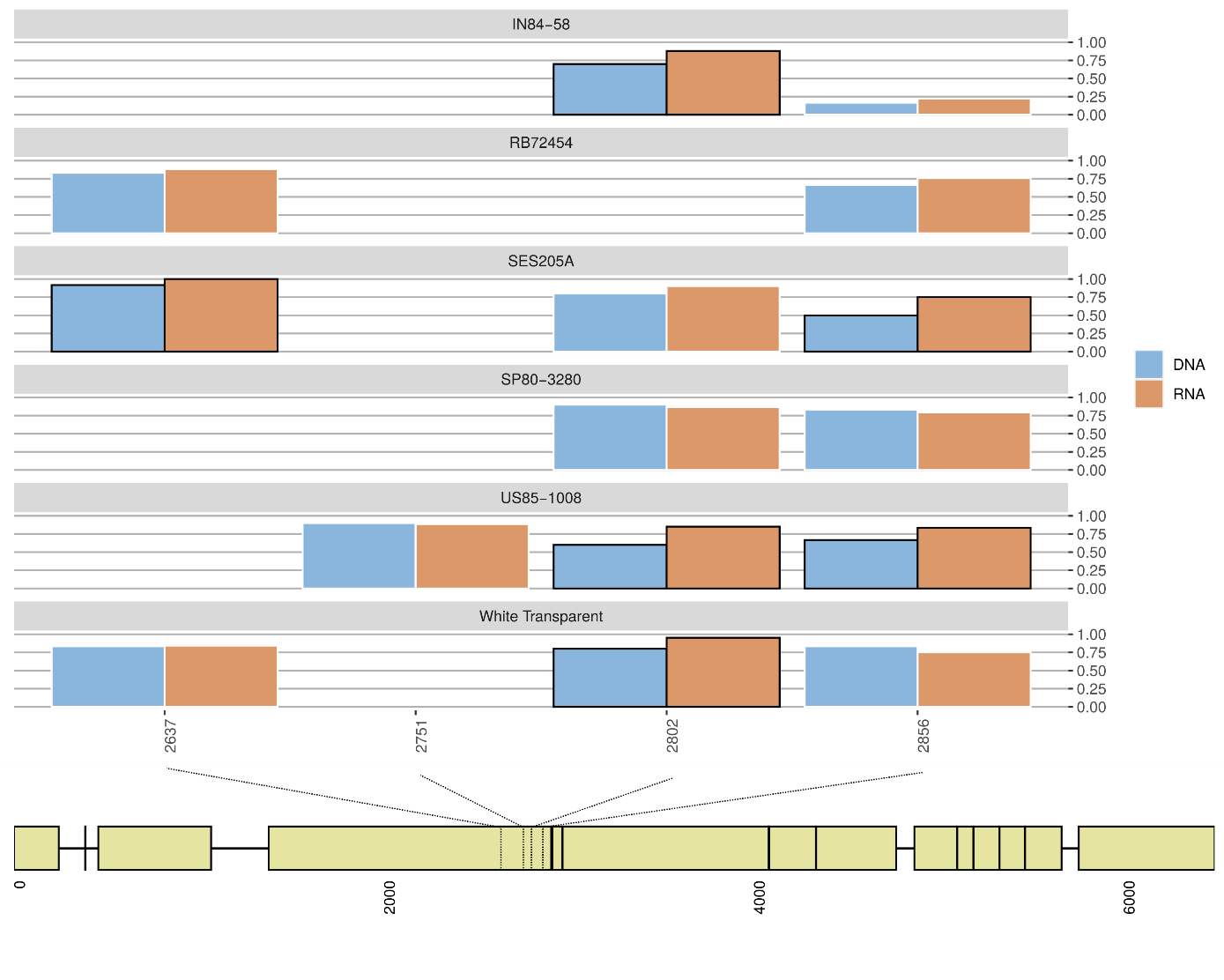

**Figure 7: Relative genomic dose and relative expression of the reference allele from SNPs identified in the gene coding for *RuBisCO large subunit-binding protein subunit alpha, chloroplastic*.** Relative genomic dose of the allele is represented by a blue bar. The expressed proportion of each allele is represented by orange bars. SNPs showing significant ASE have black borders, while those not showing ASE have white borders. The color gradient represents the average expression level of the allele. The bottom part of the plot has a schematic view of the gene showing the position of the SNPs.

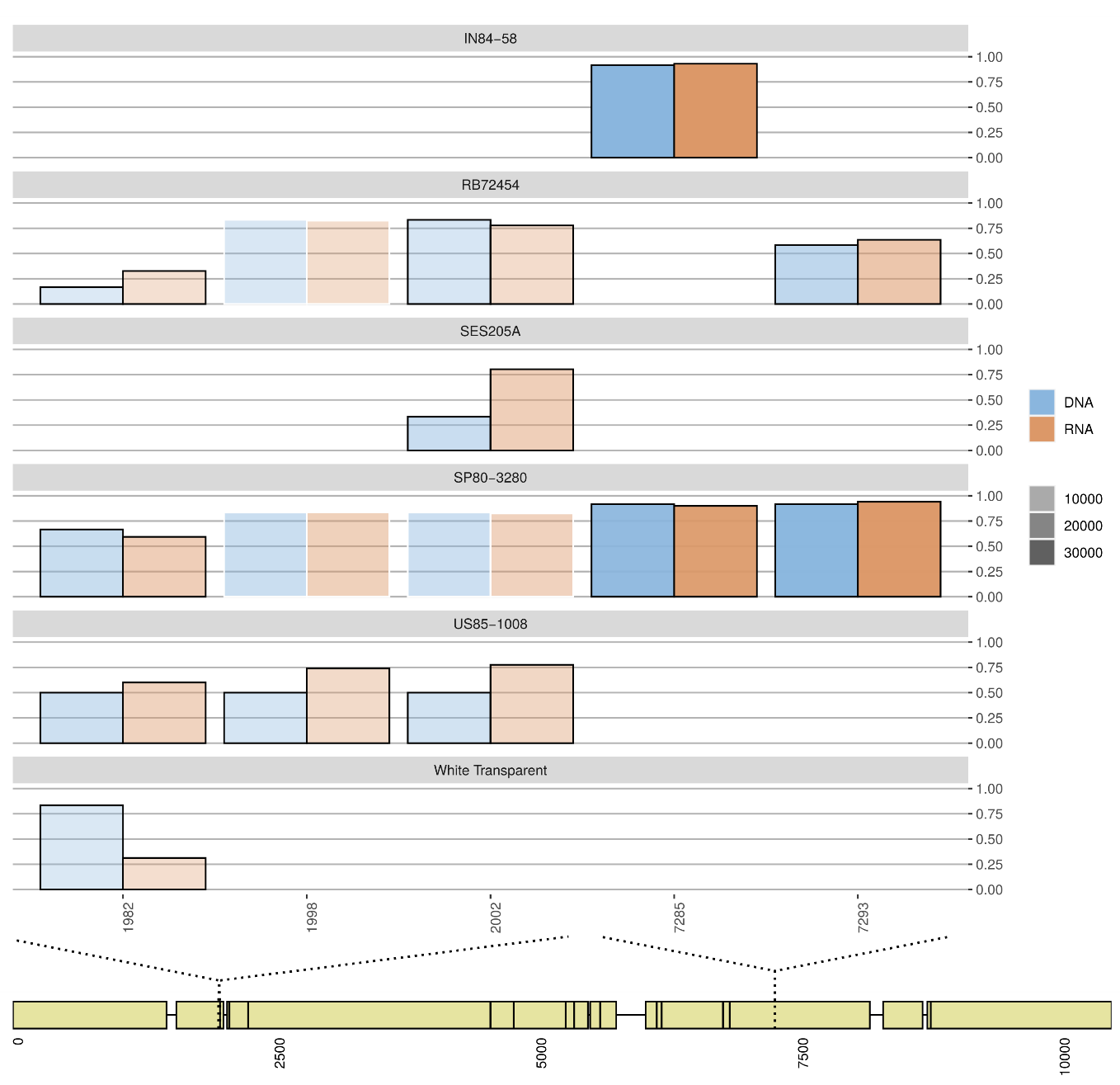

**Figure 8: Relative genomic dose and relative expression of the reference allele from SNPs identified in the gene coding for *Phosphoenolpyruvate carboxylase 3*.** Relative genomic dose of the allele is represented by a blue bar. The expressed proportion of each allele is represented by orange bars. SNPs showing significant ASE have black borders, while those not showing ASE have white borders. The color gradient represents the average expression level of the allele. The bottom part of the plot has a schematic view of the gene showing the position of the SNPs.

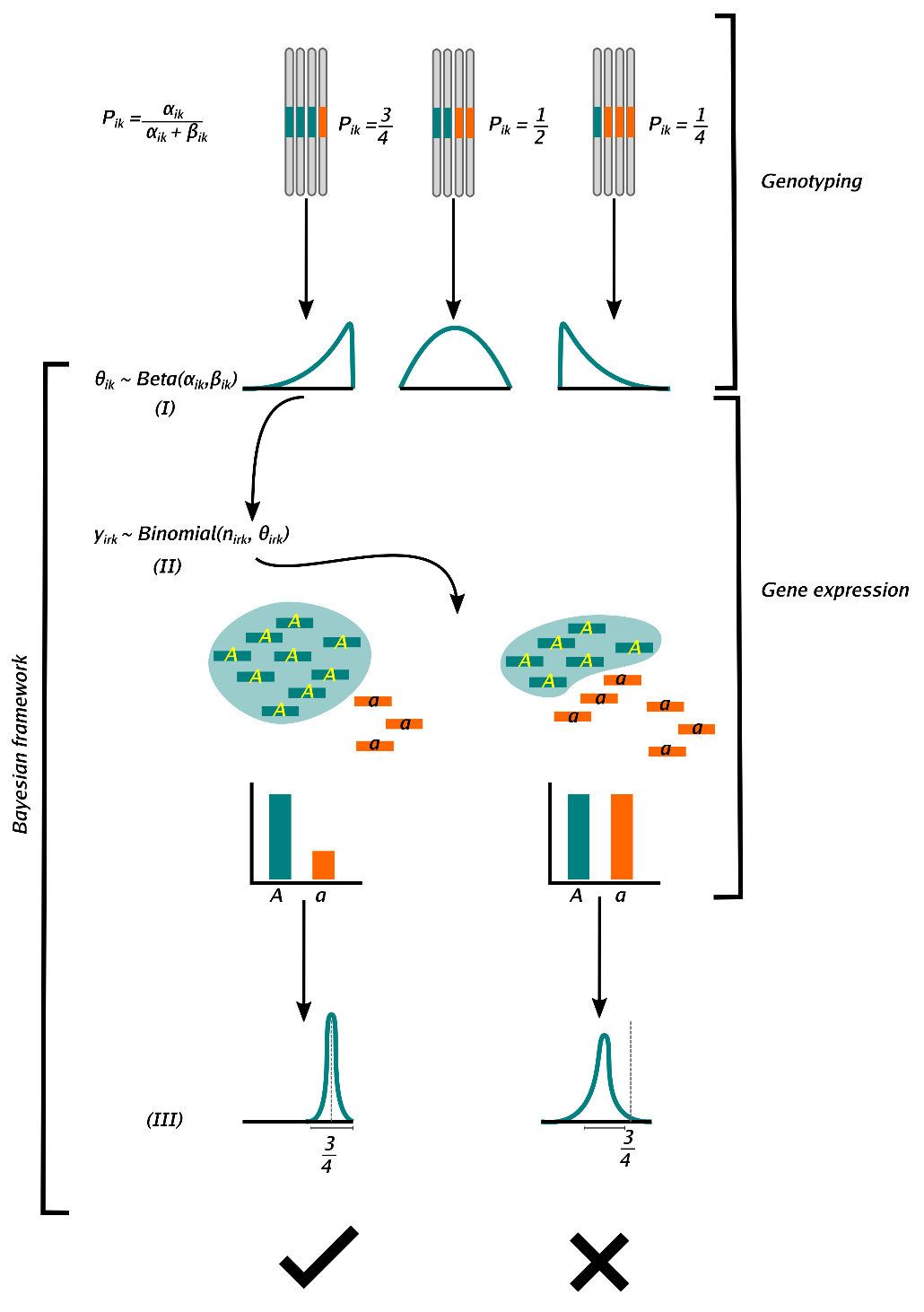

**Figure 9: Schematic view of the allele-specific expression modeling.** The genotyping section of the figure shows examples of different relative doses of the reference allele, *P_ik_*. The reference allele is colored in blue, while the alternative allele is shown in orange. In the Bayesian framework the allele doses were used as shape parameters of a beta distribution (I), which was used as the prior distribution of *θ_ik_*. From the first example of prior, we show two possible scenarios of posterior distributions. The prior was conjugated to the Binomial likelihood, which was used to model the counts associated to the reference allele (blue bars) from the total counts generated by RNA-Sequencing (II). Last, we show the posterior distribution (III), from which we tested for allele-specific expression. The check mark indicates an example of a gene with balanced expression, while the X represents a case of a gene with imbalanced expression.

##### Supplementary tables

**Table 1:** Accessions of the Brazilian Panel of Sugarcane Genotypes used in the study. For each accession we show the corresponding species, and hybrids are marked as *Saccharum* spp. The accessions were classified according to the phenotype: high or low biomass.

| Accessions | Species | Biomass group |
| --- | --- | --- |
| IN84-58 | *Saccharum spontaneum* | High |
| SES205A | *Saccharum spontaneum* | High |
| US85-1008 | *Saccharum* spp. (hybrid) | High |
| White Transparent | *Saccharum officinarum* | Low |
| RB72454 | *Saccharum* spp. (hybrid) | Low |
| SP80-3280 | *Saccharum* spp. (hybrid) | Low |

**Table 2:** Number of heterozygous SNPs in each genotype.

| Genotypes | Number of heterozygous SNPs |
| --- | --- |
| IN84-58 | 6209 |
| RB72454 | 8459 |
| SES205A | 4745 |
| SP80-3280 | 8573 |
| US85-1008 | 8901 |
| White Transparent | 8053 |

**Table 3:** Number of SNPs with significant allele-specific expression (ASE), SNPs without ASE (non-ASE), genes with ASE (ASEG) and genes without ASE (non-ASEG) per genotype.

| Genotype | Non-ASE | ASE | Non-ASEG | ASEG |
| --- | --- | --- | --- | --- |
| IN84-58 | 1370 | 804 | 625 | 467 |
| RB72454 | 1963 | 924 | 731 | 492 |
| SES205A | 1074 | 527 | 543 | 298 |
| SP80-3280 | 2115 | 987 | 819 | 517 |
| US85-1008 | 1993 | 891 | 765 | 456 |
| White Transparent | 1885 | 815 | 720 | 451 |
